# Microengineered 3D pulmonary interstitial mimetics highlight a critical role for matrix degradation in idiopathic pulmonary fibrosis

**DOI:** 10.1101/2020.06.02.129718

**Authors:** Daniel L. Matera, Katarina M. DiLillo, Makenzee R. Smith, Christopher D. Davidson, Ritika Parikh, Mohammed Said, Carole A. Wilke, Isabelle M. Lombaert, Kelly B. Arnold, Bethany B. Moore, Brendon M. Baker

**Author notes:** Corresponding Author: Brendon M. Baker, Ph.D., Assistant Professor, Department of Biomedical Engineering, University of Michigan, 2174 Lurie BME Building, 1101 Beal Avenue, Ann Arbor, MI 48109.

## Abstract

Fibrosis is often untreatable and is characterized by aberrant tissue scarring from activated myofibroblasts. Although the extracellular matrix becomes increasingly stiff and fibrous during disease progression, how these physical cues impact myofibroblast differentiation in 3D is poorly understood. Here we describe a multicomponent hydrogel that recapitulates the 3D fibrous structure hallmark to the interstitial tissue regions where idiopathic pulmonary fibrosis (IPF) initiates. In contrast to findings on 2D hydrogels, myofibroblast differentiation in 3D was inversely correlated with hydrogel stiffness, but positively correlated with matrix fiber density. Employing a multi-step bioinformatics analysis of IPF patient transcriptomes and *in vitro* pharmacologic screening, we identify matrix-metalloprotease activity to be essential for 3D but not 2D myofibroblast differentiation. Given our observation that compliant degradable 3D matrices amply support fibrogenesis, these studies demonstrate a departure from the established relationship between stiffness and myofibroblast differentiation in 2D, and provide a new 3D model for studying fibrosis.

## Introduction

Fibrosis is implicated in nearly 45% of all deaths in the developed world and plays a role in numerous pathologies including pulmonary fibrosis, cardiac disease, atherosclerosis, and cancer (*1*). In particular, interstitial lung diseases, such as idiopathic pulmonary fibrosis (IPF), are fatal and incurable with a median survival of only 2-5 years (*2*). Often described as dysregulated or incessant wound healing, fibrosis involves persistent cycles of tissue injury and deposition of extracellular matrix (ECM) by myofibroblasts (MFs). These critical cellular mediators of fibrogenesis are primarily derived from tissue resident fibroblasts. MFs drive eventual organ failure through excessive fibrous extracellular matrix deposition, force generation and tissue contraction, and eventual disruption of parenchymal tissue function (*1*). As organ transplantation remains the only curative option for late-stage disease, effective anti-fibrotic therapeutics that slow MF expansion or even reverse fibrosed tissue remain a major unmet clinical need. Undoubtedly, the limited efficacy of anti-fibrotic drugs at present underscores limitations of existing models for identifying therapeutics, the complexity of the disease, and an incomplete understanding of MF biology.

A strong correlation between lung tissue stiffening and worse patient outcomes suggests an important role for matrix mechanosensing in fibrotic disease progression (*3*). Pre-clinical models of fibrosis in mice have supported the link between tissue stiffening and disease progression. However, a precise understanding of how physical cues from the microenvironment influence MF differentiation *in vivo* is confounded by concurrent structural (eg. collagen density, laminin/elastin degradation) and biochemical (eg. matrix composition, inflammatory) changes to the microenvironment (*4*). Consequently, natural and synthetic *in vitro* tissue models have provided great utility for the study of MF mechanobiology. Seminal studies utilizing natural type I collagen gels have elucidated the role of pro-fibrotic soluble cues (e.g. TGF-β1) in promoting cell contractility, ECM compaction, and MF differentiation. However, their utility in identifying physical microenvironmental determinants of MF differentiation suffers from an intrinsic coupling of multiple biochemical and mechanical material properties (*5*). Rapid degradation kinetics (1-3 days) and resulting issues with material stability (1-2 weeks) further impede the use of natural materials for studying fibrogenic events and drug responses, which occur over weeks to months in *in vivo models* or years in patients (*6, 7*).

Synthetic hydrogels that are more resistant to cell-mediated degradation have provided a better controlled setting for long-term studies of disease-related processes (*8*). For example, synthetic hydrogel-based cell culture substrates with tunable stiffness have helped establish a paradigm for mechanosensing during MF differentiation in 2D, where compliant matrices maintain fibroblast quiescence in contrast to stiffer matrices which promote MF differentiation (*9, 10*). Extensive findings in 2D suggest a causal role for matrix mechanics (eg. stiffness) during MF differentiation *in vitro* and potentially in human disease, but these models lack the 3D nature of interstitial spaces where fibrosis originates (*11*). The interstitium surrounding alveoli is structurally composed of two key components: networks of fibrous ECM proteins (namely type I collagen fibers) and interpenetrating ground substance, an amorphous hydrogel network rich in glycosaminoglycans such as heparan sulfate proteoglycan. Mechanical cues from fibrotic ECM that promote MF differentiation may arise from changes to the collagen fiber architecture or the gel-like ground substance; indeed, whether matrix stiffness is a prerequisite for MF differentiation in 3D fibrous interstitial spaces remains unclear (*12*). Furthermore, the limited efficacy of anti-fibrotic therapies identified in pre-clinical and *in vitro* models of IPF motivates the development of 3D tissue-engineered systems with improved structural and mechanical biomimicry, relevant pharmacokinetics, and the potential to incorporate patient cells. Furthermore, recapitulating key features of the fibrotic progression in an *in vitro* setting that better approximates interstitial tissues could 1) improve our current understanding of MF mechanobiology and 2) serve as a more suitable test bed for potential anti-fibrotic therapeutics.

Accordingly, herein we describe a microengineered pulmonary interstitial matrix which recapitulates mechanical and structural features of fibrotic tissue as well as key biological events observed during IPF progression. Design parameters of these engineered microenvironments were informed by mechanical and structural characterization of fibrotic lung tissue from a bleomycin mouse model. We then investigated the influence of dimensionality, matrix crosslinking/stiffness, and fiber density on TGF-β1-induced MF differentiation in our pulmonary interstitial matrices. Surprisingly, increased hydrogel crosslinking/stiffness substantially hindered MF differentiation in 3D in contrast to findings in 2D, while fibrotic matrix architecture (i.e. high fiber density) potently promoted fibroblast proliferation and differentiation into MFs. Long-term (21 day) culture of hydrogels possessing a fibrotic architecture engendered tissue stiffening, collagen deposition, and secretion of pro-fibrotic cytokines, implicating fiber density as a potent fibrogenic cue in 3D microenvironments. Pharmacologic screening in fibrotic pulmonary interstitial matrices revealed matrix metalloprotease (MMP) activity and hydrogel remodeling as a key step during 3D fibrogenesis, but not in traditional 2D settings. To explore the clinical relevance of our findings, we leveraged a multi-step bioinformatics analysis of transcriptional profiles from 231 patients, highlighting increased MMP gene expression and enriched signaling domains associated with matrix degradation in patients with IPF. Together, these results highlight the utility of studying fibrogenesis in a physiologically relevant 3D tissue model, underscore the requirement of matrix remodeling in IPF, and establish a new platform for screening anti-fibrotic therapies.

## Results

### Evaluating the role of matrix mechanics and dimensionality on MF differentiation

To inform key design criteria for our pulmonary interstitial matrices, we began by characterizing mechanical properties of fibrotic interstitial tissue in a bleomycin-induced lung fibrosis model in mouse. Naïve C57BL/6 mice were intratracheally challenged with bleomycin to induce fibrosis, with saline treated animals maintained as a control group. After two weeks, animals were sacrificed and lung tissue was dissected out, sectioned and stained, and then mechanically tested by atomic force microscopy (AFM) nanoindentation to map the stiffness of interstitial tissue surrounding alveoli. As previously documented (*13*), bleomycin treatment corresponded to an increase in the thickness of interstitial tissue regions surrounding alveoli, a structural change which occurred alongside matrix stiffening (Fig. 1A-B); bleomycin treated lungs possessed elastic moduli nearly 5-fold greater than healthy control tissues. To generate synthetic hydrogels with elastic moduli tunable over this range, we functionalized a biocompatible and protein resistant polysaccharide, dextran, with pendant vinyl-sulfone groups amenable to peptide conjugation (termed DexVS, Fig. 1C). To permit cell-mediated proteolytic hydrogel degradation and thus spreading of encapsulated cells, we crosslinked DexVS with a bifunctional peptide (GC<u>VPMS↓MRGG</u>CG, abbreviated VPMS) primarily sensitive to MMP9 and MMP14, two MMPs implicated in fibrosis-associated matrix remodeling (*14, 15*). Tuning input VPMS crosslinker concentration yielded stable hydrogels spanning the full range of elastic moduli we measured by AFM nanoindentation of lung tissue (Fig. 1D). Additional functionalization with cell adhesive moieties (CG<u>RGD</u>S, abbreviated RGD) facilitated adhesion of primary normal human lung fibroblasts (NHLFs) (Fig. 1E).

**Figure 1:**
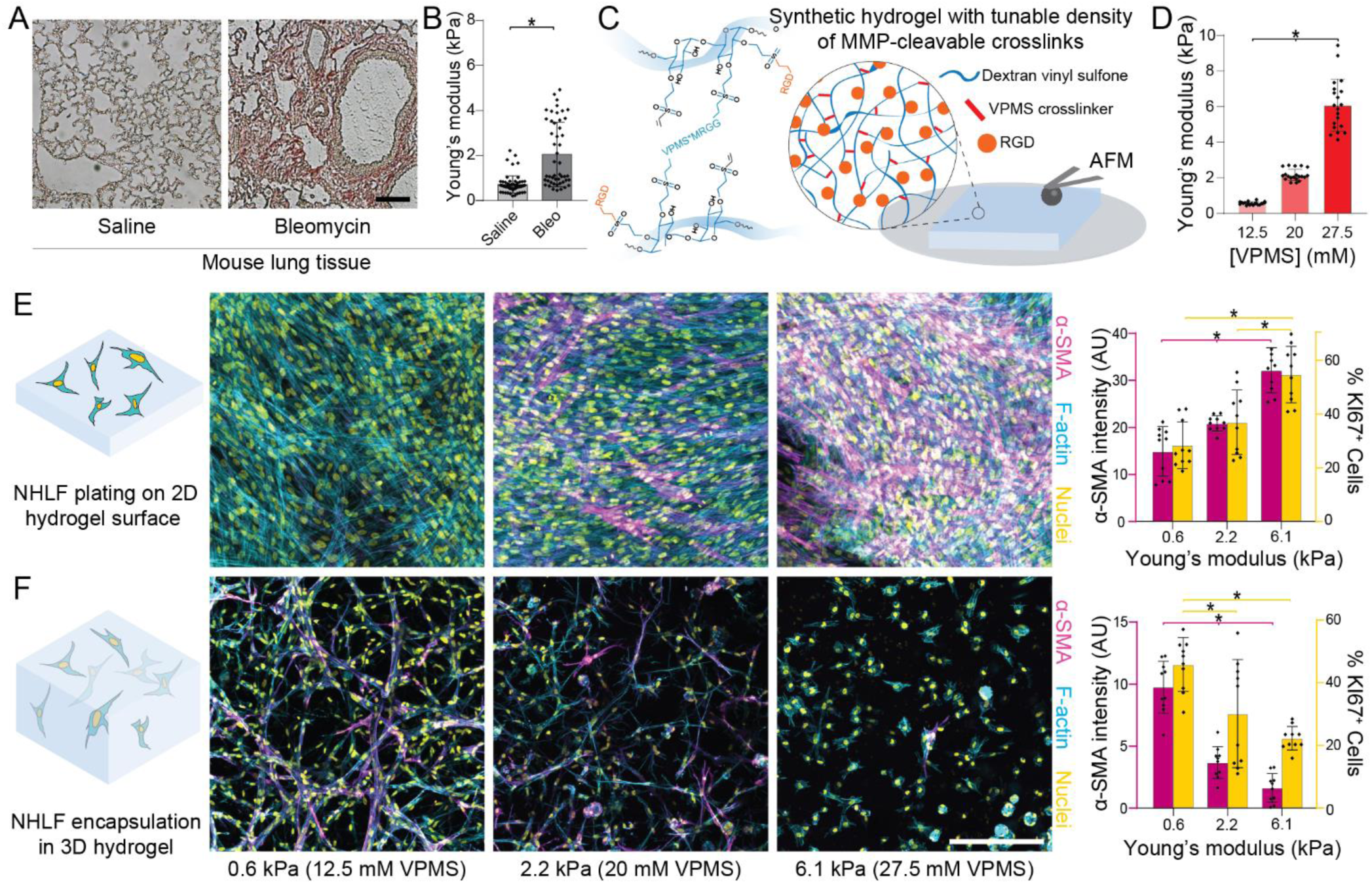
Matrix crosslinking and stiffness have opposing effects on myofibroblastic differentiation of fibroblasts plated on 2D vs. in 3D hydrogels. A) Histological preparations of healthy control and bleomycin-treated murine lung tissue (n=3 mice/group) stained for collagen by picrosirius red (scale bar: 100 µm). B) Young’s modulus of mouse lung tissue as measured by AFM nanoindentation, with data fit to the Hertz contact model to determine Young’s modulus (n=3 mice/group, n=50 indentations/group on n=9 tissue sections). C) Schematic of proteolytically sensitive, cell adhesive DexVS-VPMS bulk hydrogels. D) Young’s modulus determined by AFM nanoindentation of DexVS-VPMS hydrogels formed with different concentrations of VPMS crosslinker (n=4 samples/group, n=20 total indentations/group). E and F) Representative images of F-actin (cyan), nuclei (yellow) and α-SMA (magenta); image-based quantification of α-SMA expression and nuclear Ki67 in 2D and 3D (n=4 samples/group, n=10 fields of view/group, n>50 cells/field of view; scale bars: 200 µm). All data presented are means ± standard deviations with superimposed data points; significance determined by *p < 0.05 for all experiments.

To confirm the role of matrix mechanics on cell proliferation and MF differentiation, we seeded patient-derived NHLFs on 2D DexVS protease-sensitive hydrogel surfaces varying in VPMS crosslinker density and resulting stiffness, and stimulated cultures with TGF-β1 to promote MF differentiation. In accordance with prior literature, we observed a stiffness-dependent stepwise increase in cell proliferation (day 5) and MF differentiation (day 9) as measured by Ki67 and alpha-smooth muscle actin (α-SMA) immunofluorescence, respectively (Fig. 1E) (*10*). As the influence of matrix elasticity on MF differentiation in 3D synthetic matrices has not previously been documented, we also encapsulated NHLFs in 3D within identical DexVS hydrogels. Surprisingly, the opposing trend with respect to stiffness was noted for cells encapsulated in 3D; compliant (E=560 Pa) hydrogels which limited α-SMA expression in 2D plated cells instead exhibited the highest levels of MF differentiation in 3D (Fig. 1F). Decreasing proliferation and cell-cell contact formation as a function of increasing hydrogel stiffness were also noted in 3D matrices. Similar findings have been reported for mesenchymal stem cells (MSCs) encapsulated in hyaluronic acid matrices, where compliant gels promoted MSC proliferation and yes-associated protein (YAP) activity in 3D, yet inhibited YAP activity and proliferation in 2D (*16*). These results suggest that while stiff, crosslinked 2D surfaces promote cell spreading, proliferation, and MF differentiation, an equivalent relationship does not directly translate to 3D settings. High crosslinking and stiffness (E=6.1 kPa) in 3D matrices sterically hinder cell spreading, proliferation, and the formation of cell-cell contacts, all well-established promoters of MF differentiation (*17*).

### Microengineered pulmonary interstitial matrices to recapitulate fibrotic matrix transformation

Cell-degradable synthetic hydrogels with elastic moduli approximating that of fibrotic tissue proved non-permissive to MF differentiation in 3D. Although matrix crosslinking and densification of ground substance has previously been implicated in fibrotic tissue stiffening, remodeled collagenous architecture can also engender changes in tissue mechanics and may modulate MF development in IPF independently. To characterize the fibrous matrix architecture within healthy and fibrotic lung interstitium, we utilized second-harmonic generation (SHG) microscopy to visualize collagen microstructure in saline and bleomycin-treated lungs, respectively. Per prior literature, saline-treated lungs contained limited numbers of micron-scale (∼1 µm diameter) collagen fibers, primarily localized to the interstitial spaces supporting the alveoli (Fig. 2A) (*18*). In contrast, bleomycin-treated lungs had on average 4-fold higher overall SHG intensity, with collagen fibers localized to both an expanded interstitial region and in disrupted alveolar networks. While no difference in fiber diameter was noted with bleomycin treatment, we did observe thick (∼2-5 µm) collagen bundles containing numerous individual fibers in fibrotic lungs, potentially arising from physical remodeling by resident fibroblasts (Fig. 2A, Fig. S1). Given that typical synthetic hydrogels amenable to cell encapsulation (as in Fig. 1) lack fibrous architecture, we leveraged a previously established methodology for generating fiber-reinforced hydrogel composites (*19*). Electrospun DexVS fibers approximating the diameter of collagen fibers characterized by SHG imaging (Fig. S1) were co-encapsulated alongside NHLFs in DexVS-VPMS hydrogel matrices, yielding a 3D interpenetrating network of DexVS fibers ensconced within proteolytically cleavable DexVS hydrogel (Fig. 2B). To recapitulate the adhesive nature of collagen and fibronectin fibers within interstitial tissues, we functionalized DexVS fibers with RGD to support integrin engagement and 3D cell spreading. Importantly, while increasing the weight % of type I collagen matrices increases collagen fiber density and simultaneously increases hydrogel stiffness (Fig. S2), our synthetic matrix platform enables changes to fiber density (0.0-5.0%) without altering mechanical properties assessed by AFM nanoindentation (Fig. 2C-D), likely due to the constant weight percentage of DexVS and VPMS crosslinker within the bulk hydrogel.

**Figure 2:**
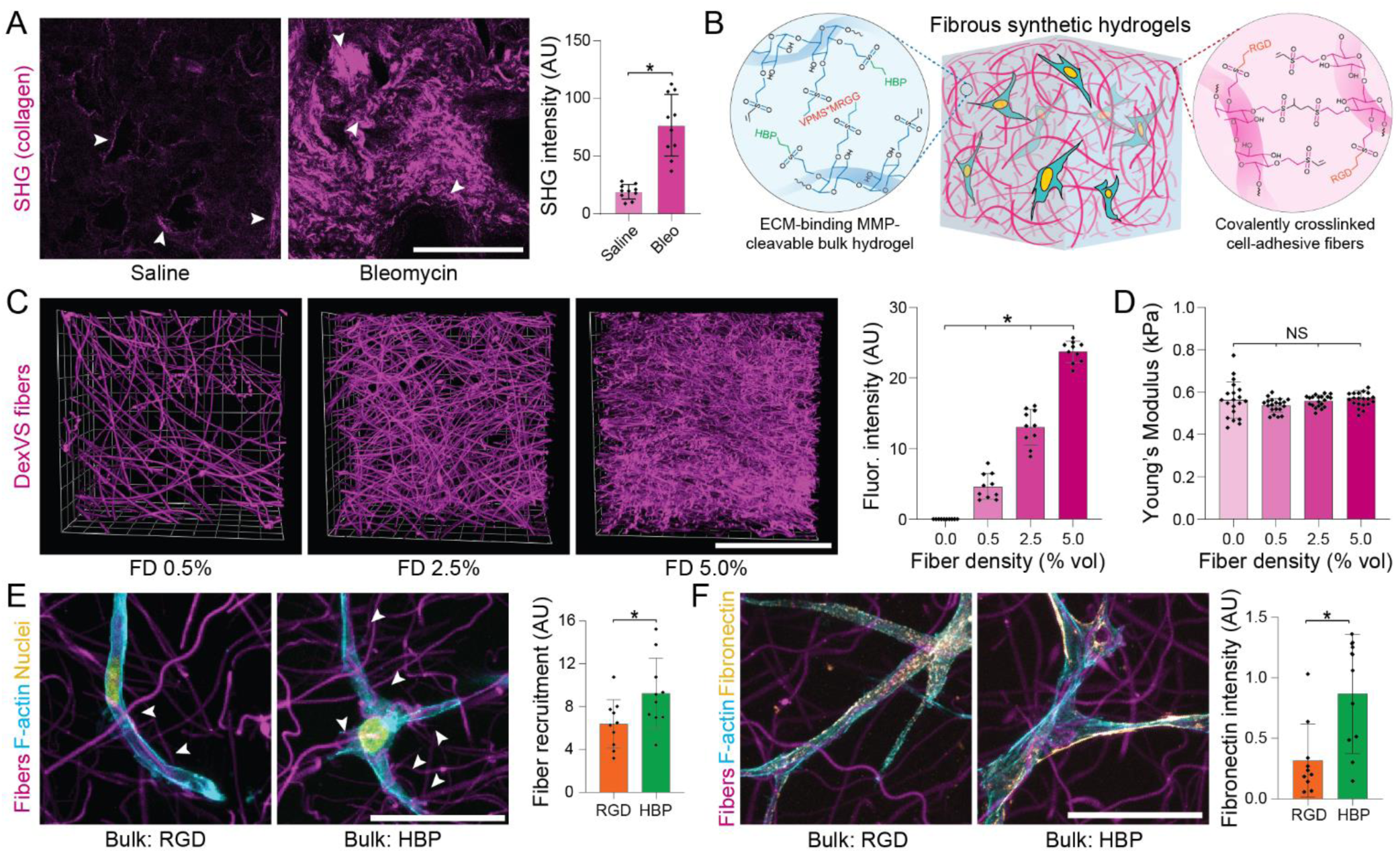
Microengineered fibrous hydrogel composites to model the lung interstitium. A) SHG imaging of collagen microstructure within healthy and bleomycin-treated lungs on day 14, with quantification of average signal intensity (arrows indicate interstitial tissue regions adjacent to alveoli; n=3 mice/group, n=10 fields of view/group; scale bar: 100 µm). B) Schematic depicting polymer crosslinking and functionalization for generating fibrous DexVS hydrogel composites to model changes in fiber density within lung interstitial tissue ECM. C) Images and intensity quantification of fluorophore-labeled fibers within composites varying in fiber density (n=4 samples/group, n=10 fields of view/group; scale bar: 100 µm). D) Young’s modulus determined by AFM nanoindentation of fibrous composites formed with different concentrations of VPMS crosslinker (n=4 samples/group, n=20 measurements/group). E) Representative high resolution images of NHLFs on day 1 in fibrous composites formed with bulk hydrogels functionalized with integrin ligand arginylglycylaspartic acid (RGD) or heparin-binding peptide (HBP) (F-actin (cyan), nuclei (yellow) and DexVS fibers (magenta); scale bar: 50 µm). Quantification of fiber recruitment as measured by contact between cells and DexVS fibers (n=10 fields of view/group, n>25 cells analyzed). F) Representative high resolution images of NHLF on day 1 fibrous composites formed with bulk hydrogels functionalized with integrin ligand RGD or heparin-binding peptide HBP (F-actin (cyan), fibronectin (yellow) and DexVS fibers (magenta); scale bar: 5 µm). Quantification of fibronectin deposition into the hydrogel matrix as measured by immunostain intensity (n=10 fields of view/group, n>25 cells analyzed). All data presented are means ± standard deviations with superimposed data points; significance determined by *p < 0.05 for all experiments.

Beyond recapitulating the multiphase structural composition of interstitial ECM, we also sought to mimic the adhesive ligand presentation and protein sequestration functions of native interstitial tissue. More specifically, the gel-like ground substance within fibrotic tissue intrinsically lacks integrin-binding moieties and is increasingly rich in heparan sulfate proteoglycans, primarily serving as a local reservoir for nascent ECM proteins, growth factors, and profibrotic cytokines. In contrast, synthetic hydrogels are often intentionally designed to have minimal interactions with secreted proteins and require uniform functionalization with a cell-adhesive ligand to support cell attachment and mechanosensing. We hypothesized that RGD-presenting fibers alone would support cell spreading (*19*) enabling the use of a non-adhesive bulk DexVS hydrogel functionalized with heparin binding peptide (HBP, CG<u>FAKLAARLYRKA</u>G). Indeed, while both RGD- and HBP-functionalized bulk DexVS gels supported cell spreading upon incorporation of RGD-presenting fibers, HBP-functionalized hydrogels encouraged matrix remodeling in the form of cell-mediated fiber recruitment (Fig. 2E) and enhanced the deposition of fibronectin fibrils into the adjacent matrix (Fig. 2F). Given the multiphase structure of lung interstitium, changes in collagen fiber density noted with fibrotic progression, and the importance of physical and biochemical matrix remodeling to fibrogenesis, we employed HBP-tethered 560 Pa DexVS-VPMS bulk hydrogels with tunable density of RGD-presenting fibers in all subsequent studies.

### Probing fibroblast-ECM interactions in a 3D tissue-engineered fibrous matrix

We next investigated whether changes in fiber density reflecting fibrosis-associated alterations to matrix architecture could influence MF differentiation in our 3D model. NHLFs were encapsulated in compliant DexVS-VPMS hydrogels ranging in fiber density (E= 560 Pa, 0.0-5.0% vol fibers). Examining cell morphology after 3 days of culture, we noted increased cell spreading and greater frequency of F-actin stress fibers and cell-cell contacts as a function of fiber density (Fig. 3A-B). As evidenced by changes in the ratio of nuclear to cytosolic YAP localization, we detected changes in mechanosensing as a function of fiber density, with the highest nuclear ratio measured in samples containing the highest fiber density examined. Given that nuclear YAP activity (a transcriptional co-activator required for downstream mechanotransduction) has been implicated as a promoter of MF differentiation, we also assayed other markers associated with fibroblast activation. With increases in fiber density, we found significant increases in cell proliferation and local fibronectin deposition (Fig. 3A-B). Luminex quantification of cytokine secretion at this time point revealed elevated secretion of inflammatory and pro-fibrotic cytokines (Fig. 3C), suggesting that matrix fibers may modulate the soluble milieu known to regulate the response to tissue damage and repair *in vivo* (*21–23*). While no α-SMA expression or collagen deposition was observed at this early time point, F-actin stress fibers, YAP activity, and fibronectin expression have been previously established as proto-MF markers *in vivo* (*24*), suggesting that physical interactions with matrix fibers primes fibroblasts for activation into MFs. Indeed, supplying the pro-fibrotic soluble factor TGF-β1 prompted increases in the expression of various pro-fibrotic YAP-target genes *(ACTA2, COL1A1, FN1, CD11, CTGF)* relative to non-fibrous (FD 0.0%) controls at day 5 (Fig. 3D). Taken together, these data suggest heightened fiber density promotes a fibrotic phenotype (Fig. 3, A to C) and gene expression (Fig. 3D), despite the absence of a stiff surrounding hydrogel.

**Figure 3:**
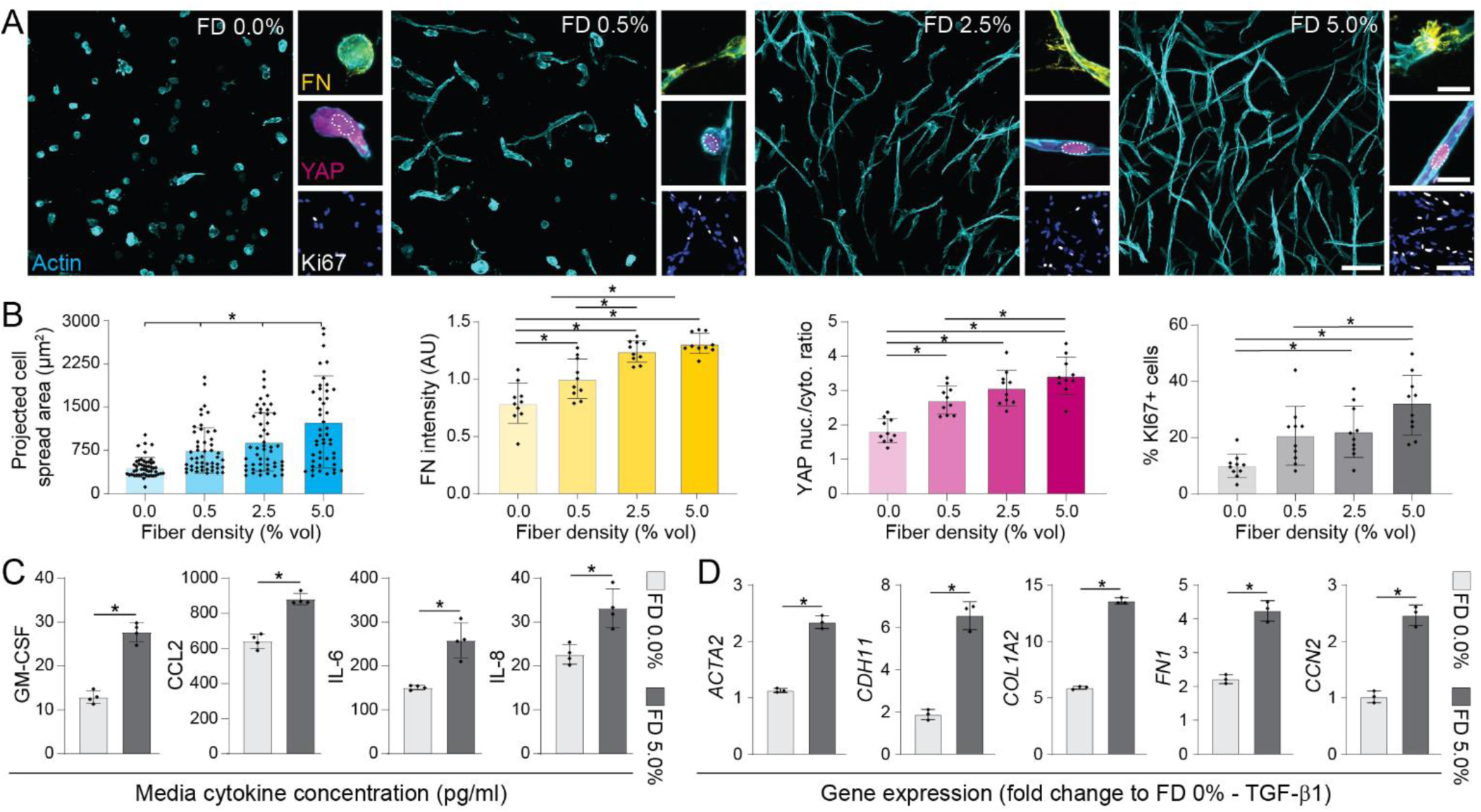
Increasing matrix fiber density in 3D fibrous hydrogels primes fibroblasts for activation into myofibroblasts. A) Immunofluorescence images of NHLFs in hydrogel composites over a range of fiber densities after 3 days of culture (F-actin (cyan), fibronectin (FN, yellow), YAP (magenta), Ki67 (white) and nuclei (blue); scale bars: F-actin 100 µm, FN 20 µm, YAP 20 µm, Ki67/nuclei 100 µm). B) Corresponding image-based quantification of cell area, deposited FN, YAP nuclear to cytosolic ratio, and % of proliferating cells (n=4 samples/group; for cell spread area analysis, n>50 cells/group; for FN, YAP, and Ki67 analyses, n=10 fields of view/group and n>25 cells/field of view). C) Cytokine secretion into culture media on day 3 (all data was normalized to background levels in control media, n=4 samples/condition). D) Expression of myofibroblast-related genes in NHLFs stimulated with TGF-β1 on day 3, either in highly fibrous (FD 5.0%) or nonfibrous (FD 0.0%) hydrogels (data presented are GAPDH-normalized fold changes relative to NHLFs within a FD 0% hydrogel lacking TGF-β1 supplementation). All data presented are means ± standard deviations with superimposed data points, significance determined by *p < 0.05 for all experiments.

### Fibrous pulmonary interstitial matrices recapitulate pathologic events observed during fibrogenesis *in vivo*

To explore whether fibrotic matrix cues in the form of heightened fiber density could promote 3D MF differentiation over longer-term culture, NHLFs were encapsulated within hydrogels varying in fiber density and maintained in media supplemented with TGF-β1 beginning on day 1. Immunofluorescent imaging and cytokine quantification were performed on days 3, 5, 7 and 9 to capture dynamic changes in cellular phenotype and secretion, respectively. No α-SMA positive stress fibers or changes in total cytokine secretion were observed on day 3 or 5. On day 7, we noted the appearance of sparse (but insignificant) α-SMA positive cells alongside increased total cytokine secretion (Fig. 4D) in FD 5.0% conditions containing TGF-β1, indicating the beginning of a potential phenotypic shift. Extensive MF differentiation and a 6-fold increase in total cytokine secretion occurred rapidly between day 7 and 9 (Fig. 4B, D-E) in the highest fiber density (FD 5.0%) condition. Importantly, despite the high proliferation within high fiber density hydrogels (Fig. 4C), MF differentiation was not evident in samples lacking exogenous TGF-β1 supplementation. Moreover, 3D MF differentiation was also absent in TGF-β1 supplemented conditions that lacked fibrous architecture, indicating a requirement for both soluble and physical fibrogenic cues in 3D. Furthermore, inhibiting integrin engagement by incorporating fibers lacking RGD also abrogated MF differentiation and proliferation despite the presence of TGF-β1 (Fig. 4A-B), suggesting that a fibrotic matrix architecture drives α-SMA expression primarily through integrin engagement and downstream mechanosensing pathways. These results were replicated with primary human dermal fibroblasts (NHDF) and mammary fibroblasts (NHMF), where similar trends with α-SMA expression as a function of fiber density were observed (Fig. S3). Interestingly, while high fiber density promoted proliferation in NHDFs, NHMFs underwent MF differentiation in the absence of higher proliferation rates, demonstrating intrinsic differences between cell populations originating from different tissues. Nevertheless, these results suggest that fibrotic matrix architecture may be promoting MF differentiation in other pathologies, namely dermal scarring in systemic sclerosis and desmoplasia in breast cancer.

**Figure 4:**
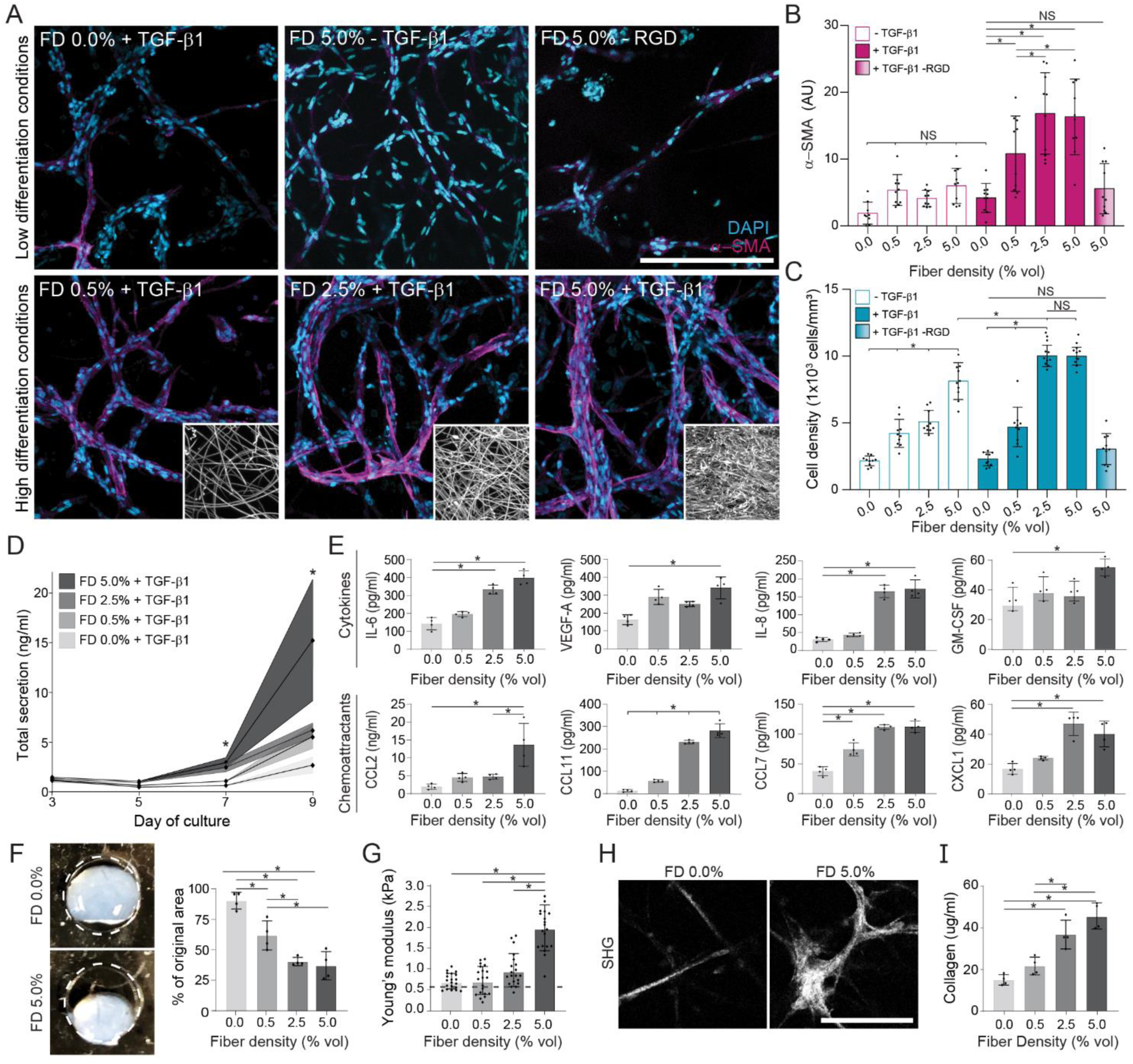
Profibrotic soluble and physical cues promote MF differentiation in 3D and initiate a progression of fibrosis-associated changes over long-term culture. A) Representative immunofluorescence images of NHLFs in microenvironmental conditions leading to low or high MF differentiation after 9 days in culture (α-SMA (magenta) and nuclei (cyan); n=4 samples/group, n=10 fields of view/group and n>50 cells/field of view; scale bar: 200 µm), with corresponding image-based quantification in (B) and (C). Insets depict representative fiber densities. D) Measurement of total cytokine secretion over time as a function of fiber density (n=4 samples/condition, * indicates significant differences between FD 5.0% and all other groups at a given time point). E) Secretion of specific cytokines and chemoattractants as a function of fiber density on day 9 (n=4 samples/condition). F) Representative images and quantification of tissue contraction within day 14 fibroblast-laden hydrogels of varying fiber density (n=4 samples/group, dashed line indicates initial diameter of 5 mm). G) AFM measurements of day 14 fibroblast-laden hydrogels of varying fiber density (n=20 measurements from n=4 samples/group). Dashed line indicates original hydrogel stiffness. H) SHG images of fibrous collagen within fibroblast laden hydrogels after 21 days of culture in media supplemented with ascorbic acid (scale bar: 100 µm). I) Measurement of total collagen content within digested DexVS hydrogels at day 21 as measured by biochemical assay (n=4 samples/group). All data presented are means ± standard deviations with superimposed data points; significance determined by *p < 0.05 for all experiments.

While proliferation and α-SMA expression are accepted markers of activated fibroblasts, fibrotic lesions contribute to patient mortality through airway inflammation, collagen secretion, tissue contraction, and lung stiffening – pathogenic events which hinder the physical process of respiration (*25*). Luminex screening of 41 cytokines and chemokines within hydrogel supernatant revealed elevated total cytokine secretion as a function of fiber density over time (Fig. 4D), many of which were soluble mediators known to regulate airway inflammation (Fig. 4E) (*21*). Numerous other cytokines were additionally secreted at day 9 but did not change as a function of fiber density (Fig. S4). By generating free-floating hydrogels which allow for contraction over time, we also examined macroscale changes in tissue geometry. Consistent with the influence of fiber density on α-SMA expression, hydrogels containing high fiber densities underwent greater hydrogel contraction compared to non-fibrous or low fiber density conditions (Fig. 4F). Day 14 fibrotic tissues (FD 5.0%) were also 4-fold stiffer (2.0 vs. 0.5 kPa) as measured by AFM nanoindentation (Fig. 4G) compared to conditions which yielded low rates of MF differentiation in shorter term studies (ie. FD 0.0 or FD 0.5% in Fig. 4A-B). When media was supplemented with ascorbic acid to permit procollagen hydroxylation, collagen deposition into surrounding matrix was evident by SHG microscopy by day 21 in high fiber density hydrogels (Fig. 4H) as compared to nonfibrous controls. Further biochemical analysis of hydrogel collagen content confirmed a stepwise increase in collagen production as a function of fiber density (Fig. 4I). Taken together, these findings demonstrate a clear influence of fiber density on MF differentiation and phenotype in 3D, and furthermore suggest that this *in vitro* model recapitulates key pathogenic events associated with the progression of fibrosis *in vivo.*

### Therapeutic screening in fibrotic matrices alongside transcriptional profiling of IPF patients highlight the requirement for matrix remodeling during MF differentiation and in clinical fibrosis

Having established microenvironmental cues that promote robust 3D MF differentiation, we next evaluated the potential of our fibrous hydrogel model for use as an anti-fibrotic drug screening platform. Nintedanib (NTN), a broad-spectrum receptor tyrosine kinase (RTK) inhibitor, and Pirfenidone (PFD), an inhibitor of the MAPK/NF-κB pathway, were selected due to their recent FDA approval for use in patients with IPF (*26*). We also included dimethylformamide (DMF), an inhibitor of the YAP/TAZ pathway clinically approved for treatment of systemic sclerosis, and Marimastat (MMS), a broad-spectrum MMP inhibitor which has shown efficacy in murine pre-clinical models of fibrosis (*27, 28*). We generated fibrotic matrices (560 Pa DexVS-VPMS-HBP bulk hydrogels containing 5.0% vol DexVS-RGD fibers) which elicited the highest levels of MF differentiation, matrix contraction, and collagen secretion in our previous studies (Fig. 4). As a comparison to the current standard for high throughput compound screening, we also seeded identical numbers of NHLFs on 2D tissue culture plastic (TCP) in parallel. Cultures were stimulated with TGF-β1 on day 1, and pharmacologic treatments were added on day 3, following extensive fibroblast spreading, cell-cell junction formation, and proliferation (Fig. 3A).

As in our earlier studies, TGF-β1 supplementation promoted proliferation and α-SMA expression within 3D constructs as well as on on rigid tissue culture plastic (Fig. 5a). Interestingly, NTN and PFD had differential effects on NHLFs depending on culture format; NHLFs on 2D TCP were resistant to PFD/NTN treatment with no difference in proliferation or α-SMA expression relative to vehicle controls, whereas modest but significant decreases in α-SMA expression (PFD, NTN) and proliferation (NTN) were detected in 3D (Fig. 5, A to E). Combined treatment with PFD and NTN provided an anti-fibrotic effect only in fibrotic matrices, supporting ongoing clinical studies exploring their use as a combinatorial therapy (ClinicalTrials.gov identifier NCT03939520). DMF abrogated cell proliferation and α-SMA expression across all conditions, suggesting that inhibition of downstream mechanosensing inhibits MF differentiation in both 2D and 3D contexts in support of the general requirement for mechanosensing during MF differentiation independent of culture substrate (*10*). Indeed, inhibition of YAP activity *in vivo* has been shown to mitigate fibrosis and may be an advantageous therapeutic target (*20*). Blockade of MMP activity via MMS treatment proved ineffectual in reducing α-SMA expression or proliferation on 2D TCP, but surprisingly fully abrogated the proliferation and differentiation response in 3D fibrotic matrices (Fig. 5, A to E). Given the role of protease activity in tissue remodeling *in vivo* (*28*) and in cellular outgrowth within 3D hydrogels (*16, 29*), our data suggest that degradative matrix remodeling is a requirement for MF differentiation in 3D, but not in more simplified 2D settings. To summarize, multiple anti-fibrotic agents (PFD, NTN, DMF, MMS) demonstrating efficacy in clinical literature elicited an anti-fibrotic effect in our engineered fibrotic pulmonary interstitial matrices, but not in the 2D TCP contexts traditionally used for compound screening.

**Figure 5:**
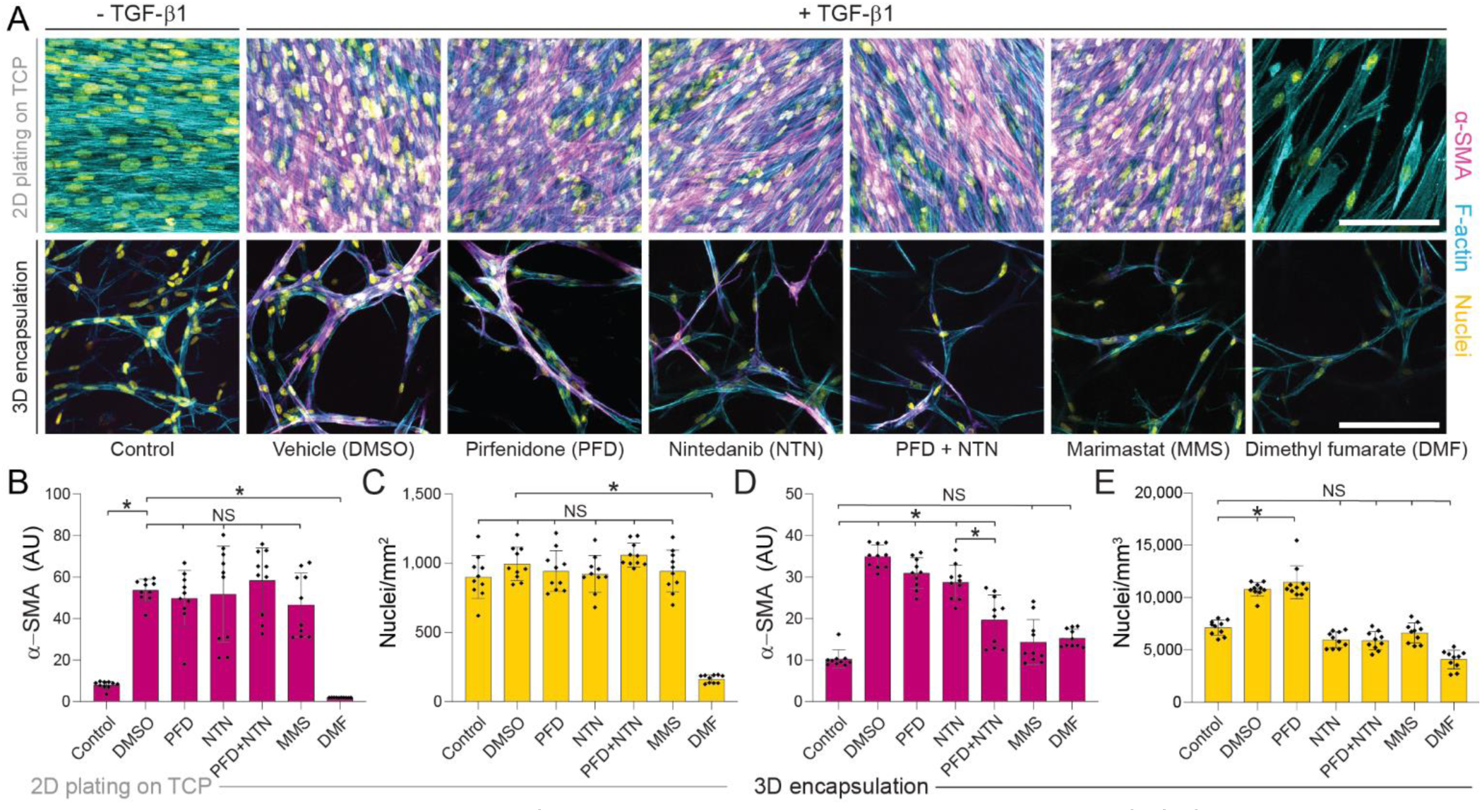
Pharmacologic treatment of NHLFs plated on tissue culture plastic (TCP) in 2D compared to encapsulated within 3D fibrous hydrogels reveals differential drug responses. A) Representative confocal images stained for α-SMA (magenta), F-actin (cyan), and nuclei (yellow) of NHLFs after 9 days of culture on TCP (top row) or 3D fibrotic matrices (bottom row) with pharmacologic treatment indicated from days 3 to 9 (scale bar: 100 µm). Imaged regions were selected to maximize the # of α-SMA+ cells/field of view within each sample. B) Quantification of α-SMA and C) total cell count within 2D NHLF cultures. D) Quantification of α-SMA and E) total cell count within 3D fibrotic matrices (n=4 samples/group, n=10 fields of view/group and n>50 cells/field of view). All data presented are means ± standard deviations with superimposed data points; significance determined by *p < 0.05 for all experiments.

As the protease inhibitor MMS fully ablated TGF-β1 induced α-SMA expression and proliferation in our 3D fibrotic matrices, we leveraged bioinformatics methodologies to investigate the role of matrix proteases in IPF patients on a network (Reactome) and protein (STRING) basis. Differential expression analysis of microarray data within the NCBI GEO (dataset # GSE47460) was used to generate an uncurated/unbiased dataset composed of the top 1000 differentially regulated genes in IPF, revealing *MMP1* as the most upregulated gene in IPF patients, with other matrix proteases (*MMP1, MMP3, MMP7, MMP9, MMP10, MMP11, MMP12*) and matrix remodeling proteins (*COL1A2, LOX, ACAN, DCN, HS6ST2*) similarly upregulated (Fig. 6B, Table S1, Data File S1). To focus on genes associated with MF differentiation for subsequent analyses, we performed GO term enrichment (via GEO2R) to compile a curated dataset containing 188 key genes associated with MF differentiation (Data File S1) and utilized Reactome and STRING analyses to investigate network signaling within both the uncurated and curated datasets. Analyses revealed 103 (uncurated) and 89 (curated) enriched signaling pathways in IPF (Data File S1). Interestingly, the top 3/5 (uncurated) and 5/5 (curated) significantly enriched pathways in IPF involved matrix degradation and remodeling (Fig. 6C). Subsequent STRING protein-protein interaction analysis of datasets revealed that top signaling nodes were MMPs (uncurated: *MMP1, MMP3*; Fig. 6D), fibrous collagens (uncurated: *COL1A2, COL3A1*), or cytokines (curated: *IL6, VEGFA, IL1B, IGF1*; Fig. 6D) known to increase MMP expression in fibroblasts (*30–33*). These results emphasize the interdependence between MMP activity and systems-level pathogenic signaling in IPF, and in combination with our 3D drug screening results, highlight fibroblast-specific protease activity as a potential therapeutic target. Furthermore, given that protease inhibition had no effect on MF differentiation in 2D culture, these data also support the growing sentiment that simplified 2D screening models may be masking the identification of potentially viable anti-fibrotics.

**Figure 6:**
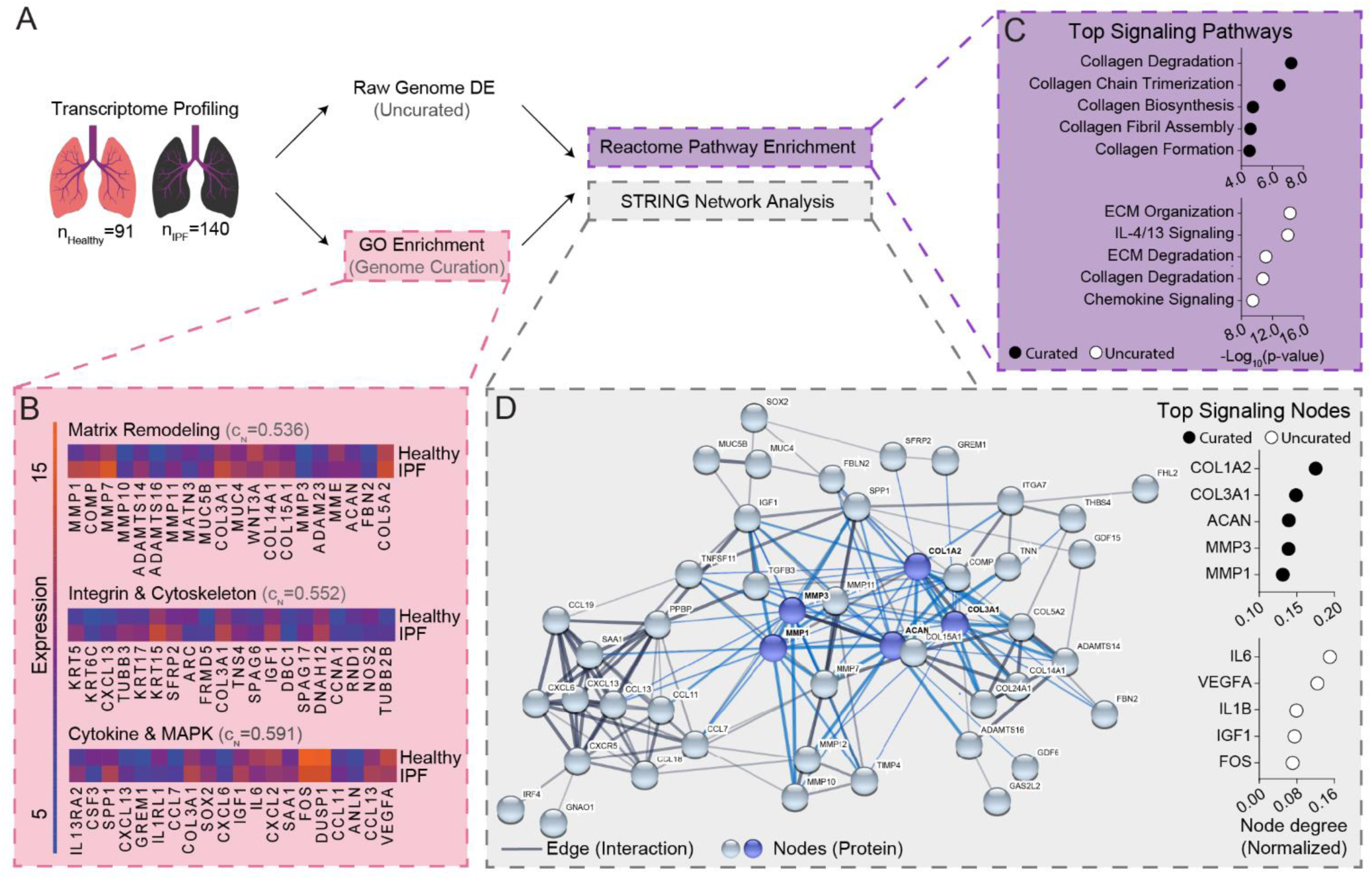
Bioinformatics analysis of IPF patient transcriptomes reveals matrix remodeling as a key signaling node in human disease. A) Schematic representation of bioinformatics workflow: whole-genome transcriptomes from 91 healthy and 140 patients with lung fibrosis were fetched from the NCBI GEO. Differential expression analysis was utilized to assemble an uncurated list of the top 1000 differentially expressed genes. GO enrichment of choice biological pathways was utilized to assemble a curated list of genes associated with MF differentiation. Datasets were fed through a prior-knowledge based analysis pipeline to identify enriched signaling pathways (Reactome) and key protein signaling nodes (STRING) within IPF patients. B) Heatmaps of the top 20 differentially expressed genes within specified GO categories which were manually selected for curated analysis. C_N_ values indicate a high degree of interaction between proteins selected for curated analysis. C) Summary of the top 5 significantly enriched pathways in the curated and uncurated gene set. D) Representative STRING diagram depicting protein interactions within the curated dataset, with summary of the top 5 signaling nodes in the uncurated and curated gene set. Blue nodes and edges represent interactions within the top 5 signaling nodes for the curated dataset.

## Discussion

Despite fibrosis widely contributing to mortality worldwide, inadequate understanding of fibrotic disease pathogenesis has limited the development of efficacious therapies (*11*). Pre-clinical studies *in vivo*, while indispensable, often fail to translate to clinical settings as evidenced by the failure of ∼90% of drugs identified in animal studies (*34*). Additionally, limitations in current technologies (i.e. the embryonic lethality of many genetic ECM knockouts and the limited resolution/imaging depth of intravital microscopy) have hindered the application of pre-clinical *in vivo* models for the study of cell-ECM interactions which underlie fibrogenesis (*35*). In contrast, existing *in vitro* models utilize patient-derived cells are affordable, scalable, and amenable to microscopy, but often fail to recapitulate the complex 3D matrix structure hallmark to the interstitial tissue regions where fibrotic diseases such as IPF originate. We leveraged electrospinning and bio-orthogonal chemistries to engineer novel pulmonary interstitial matrices that are 3D and possess fibrous architecture with biomimetic ligand presentation. In the presence of a pro-fibrotic soluble factors, these settings reproduce hallmarks of fibrosis at cellular and tissue levels (Fig. 2-4). Examining the influence of physical microenvironmental cues (crosslinking/stiffness and fiber density) on MF differentiation, we find that crosslinking/stiffness has opposing effects on MF differentiation in 2D vs. 3D (Fig. 1), and that incorporation of a fibrous architecture in 3D is a prerequisite to MF differentiation (Fig. 4). Furthermore, supported by the importance of protease signaling in IPF (Fig. 6), we performed proof-of-concept pharmacologic screening within our 3D fibrotic matrices (Fig. 5) and highlighted enhanced biomimicry as compared to traditional 2D drug screening substrates where matrix remodeling appears to be dispensable for MF differentiation.

While tunable synthetic hydrogels have identified mechanosensing pathways critical to MF differentiation in 2D, these observations have yet to be translated to 3D, fibrous settings relevant to the interstitial spaces where fibrosis originates. Given that late-stage IPF progresses in the absence of external tissue damage, current dogma implicates fibrotic matrix stiffness as the continual driver of MF differentiation *in vivo* (*9, 10, 36*). While we cannot disregard this hypothesis, our work elucidates a contrasting MMP-dependent mechanism at play in 3D, whereby a compliant, degradable, and fibrous matrix architecture supports MF differentiation, with matrix contraction and stiffening occurring downstream of α-SMA expression, nearly a week later. Given numerous 2D studies indicating matrix stiffness as a driver of MF differentiation, the finding that a compliant matrix promotes MF differentiation may appear counterintuitive (*9, 10*). However, MF accumulation has been documented prior to tissue stiffening in human disease (*3*), and a recent phase 2 clinical trial (ClinicalTrials.gov Identifier: NCT01769196) targeting the LOX pathway (the family of enzymes responsible for matrix stiffening *in vivo*) failed to prevent disease progression in IPF patients and was terminated due to lack of efficacy (*37*). Furthermore, compelling recent work by Fiore et. al. combined immunohistochemistry with high-resolution AFM to characterize human IPF tissue mechanics, and found that regions of active fibrogenesis were highly fibrous but possessed a similar Young’s modulus as healthy tissue (*3*). In concert with our *in vitro* data, these findings suggest that MF differentiation is possible within soft provisional ECM *in vivo*, and that the initiation of fibrogenesis may not be dependent on heightened tissue stiffness, so long as matrix fibers and appropriate soluble cues (e.g. TGF-β1) are present.

Consequently, understanding the source of pro-fibrotic soluble cues *in vivo* is of critical importance when identifying therapeutic targets for IPF. Luminex screening of supernatant from 3D fibrotic matrices revealed 6-fold increases in cytokine secretion during fibrogenesis, the majority of which were potent inflammatory factors (eg. GM-CSF, IL-6, IL-8, VEGF-A) and chemoattractants (eg. CCL2, CCL7, CCL11, CXCL1) (Fig. 4E). Furthermore, IL-6 and VEGF-A were found to be major signaling nodes in IPF patients (Fig. 6D). While not typically regarded as an immunomodulatory cell population, these findings suggest MFs may maintain localized inflammation to support continual fibrogenesis. Mitogens such as IL-6 and IL-8 promote endothelial- and epithelial-to-mesenchymal transition, a process that gives rise to matrix-producing MF-like cells in IPF (*38*). CCR2 (CCL2, CCL7) and CXCR1 (CXCL1, IL-8) ligation facilitates macrophage chemotaxis, potentially leading to a sustained influx of TGF-β1 producing cells in IPF, and glycoproteins such as GM-CSF inhibit caspase activity in mononuclear cells, potentially preventing apoptotic events required for the resolution of wound repair and return to homeostasis (*21, 39*). Additionally, secretion of nearly all cytokines were increased as a function of fiber density, highlighting a potential feed-forward loop distinct from canonical TGF-β1 signaling. Further model development (i.e. co-culture platforms) will be required to examine these hypotheses and the role of MF-derived cytokines in persistent inflammation and fibrosis.

In addition to documenting the role of fibrotic matrix architecture in 3D fibrogenesis, we demonstrate proof-of-concept pharmacologic screening within our synthetic pulmonary interstitial matrices and highlight their improved relevance to human disease. Prior work *in vitro* has documented profound reductions in MF differentiation after treatment with clinically approved anti-fibrotics (PFD, NTN), whereas in the clinic, PFD and NTN impede disease progression but are far from curative (*4, 26, 40, 41*). Interestingly, PFD or NTN had insignificant effects in 2D settings in our hands and only modest effects in 3D (Fig. 5). One reason for this discrepancy may be the use of supraphysiologic PFD and NTN concentrations in previous *in vitro* studies, whereas we selected dosages based upon plasma concentrations in IPF patients (*42*). Differences in pharmacokinetics and cell metabolism between 2D and 3D tissue constructs likely also play a role. Furthermore, as evidenced by the preventative effect of the protease inhibitor Marimastat in 3D hydrogels but not 2D settings (Fig. 5), pharmacologics which influence matrix degradation and remodeling are likely to have a minimized effect in 2D settings due to the less dynamic nature of TCP and flat hydrogels (*43*). Indeed, NTN and PFD have been shown to influence protease activity and matrix remodeling *in vivo* (*15*), and may be mediating their effects within fibrotic matrices through modulation of ECM remodeling. Given the identification of numerous potential anti-fibrotic agents (microRNA, TGF-β1 inhibitors, IL-4, IL-13 neutralizing antibodies, integrin blockers) in pre-clinical models, application of the system described here could elucidate how choice pharmacologics impact MF differentiation and matrix remodeling processes which are difficult to recapitulate in 2D culture. Further development of our interstitial matrices as an arrayed microtissue platform, as has been elegantly implemented with collagen matrices (*40*), is a critical next step to moving this technology towards high-throughput screening applications.

In summary, we designed a tunable 3D and fibrous hydrogel model which recapitulates dynamic physical (eg. stiffening, contraction) and biochemical (eg. secretion of fibronectin, collagen, and cytokines) alterations to the microenvironment observed during the progression of IPF. Implementation of our model allowed us to establish a developing mechanism for MF differentiation in 3D compliant environments, whereby cell spreading upon matrix fibers drives YAP activity, cytokine release, and proteolysis-dependent MF differentiation. Furthermore, we leveraged bioinformatics techniques to explore protease signaling in clinical IPF, and in concert with our therapeutic screening data, establish a strong role for matrix degradation during IPF pathogenesis and in 3D MF differentiation, respectively. Consequently, these results highlight critical design parameters (3D degradability, matrix architecture) frequently overlooked in established synthetic models of MF differentiation. Future work incorporating macrophages, endothelial cells and epithelial cells may expand current understanding of how developing MF populations influence otherwise homeostatic cells, and how matrix remodeling influences paracrine signaling networks and corresponding drug response. Given the low translation rate of drugs identified in high-throughput screening assays, we show that the application and development of engineered biomimetics, in combination with pre-clinical models, can improve drug discovery and pathophysiological understanding.

## Materials and Methods

### Reagents

All reagents were purchased from Sigma Aldrich and used as received, unless otherwise stated.

### Synthesis and characterization of modified dextran

<u>Dextran vinyl sulfone (DexVS)</u>: A previously described protocol for vinyl sulfonating polysaccharides was adapted for use with linear high MW dextran (MW 86,000 Da, MP Biomedicals, Santa Ana, CA) (*19*). Briefly, pure divinyl sulfone (12.5 ml, Fisher Scientific, Hampton, NH) was added to a sodium hydroxide solution (0.1 M, 250 mL) containing dextran (5 g). This reaction was carried out at 1500 RPM for 3.5 minutes, after which the reaction was terminated by adjusting the pH to 5.0 via the addition of hydrochloric acid. A lower functionalization of DexVS was utilized for hydrogels, where the volume of divinyl sulfone reagent was reduced to 3.875 ml. All reaction products were dialyzed for 5 days against Milli-Q ultrapure water, with two water exchanges daily, and then lyophilized for 3 days to obtain the pure product. Functionalization of DexVS was characterized by ^1^H –NMR spectroscopy in D_2_O and was calculated as the ratio of the proton integral (6.91 ppm) and the anomeric proton of the glucopyranosyl ring (5.166 and 4.923 ppm), here a vinylsulfone/dextran repeat unit ratio of 0.376 and 0.156 was determined for electrospinning and hydrogel DexVS polymers, respectively.

### Fiber segment fabrication

DexVS was dissolved at 0.6 g ml^-1^ in a 1:1 mixture of Milli-Q ultrapure water and dimethylformamide with 0.015% Irgacure 2959 photoinitiator. Methacrylated rhodamine (0.5 mM; Polysciences, Inc., Warrington, PA) was incorporated into the electrospinning solution to fluorescently visualize fibers under 555 laser. This polymer solution was utilized for electrospinning within an environment-controlled glovebox held at 21 °C and 30% relative humidity. Electrospinning was performed at a flow rate of 0.3 ml h^-1^, gap distance of 5 cm, and voltage of -10.0 kV onto a grounded collecting surface attached to a linear actuator. Fiber layers were collected on glass slabs and primary crosslinked under ultraviolet light (100mW cm^-2^) and then secondary crosslinked (100 mW cm^-2^) in a 1mg mL^-1^ Irgacure 2959 solution. After polymerization, fiber segments were resuspended in a known volume of PBS (typically 3 ml). The total volume of fibers was then calculated via a conservation of volume equation: total resulting solution volume = volume of fibers + volume of PBS (3 ml). After calculating total fiber volume, solutions were re-centrifuged, supernatant was removed, and fiber pellets were resuspended to create a 10 vol% fiber solution, which were then aliquoted and stored at 4° C. To support cell adhesion, 2.0 mM RGD was coupled to vinyl sulfone groups along the DexVS backbone via Michael-type addition chemistry for 30 min, followed by quenching of excess VS groups in a 300 mM cysteine solution for 30 minutes.

### Hydrogel formation

DexVS gels were formed via a thiol-ene click reaction at 3.3% w/v (pH 7.4, 37°C, 45 min) with VPMS crosslinker (12.5, 20, 27.5 mM) (GCRDVPMSMRGGDRCG, Genscript, George Town, KY) in the presence of varying amounts of argininylglycylaspartic acid (RGD, CGRGDS, 2.0 mM, Genscript, George Town, KY), heparin-binding peptide (HBP, GCGAFAKLAARLYRKA, 1.0 mM, Genscript, George Town, KY), and fiber segments (0.0-5.0% v/v). For experiments comparing hydrogels of varying ligand type (1 mM HBP vs. 2 mM RGD) cysteine was added to precursor solutions to maintain a final vinyl sulfone concentration of 60 mM. All hydrogel and peptide precursor solutions were made in PBS containing 50 mM HEPES. To create fibrous hydrogels, a defined stock solution (10% v/v) of suspended fibers in PBS/HEPES was mixed into hydrogel precursor solutions prior to gelation. Via controlling the dilution of the fiber suspension, fiber density was readily tuned within the hydrogel at a constant hydrogel weight percentage. For gel contraction experiments, DexVS was polymerized within a 5 mm diameter PDMS gasket to ensure consistent hydrogel area on day 0.

### Cell culture and biological reagents

Normal Human Lung Fibroblasts (NHLFs, University of Michigan Central Biorepository), Normal Human Dermal Fibroblasts (NHDF, Lonza, Basel, Switzerland), and Normal Human Mammary Fibroblasts (NHMF, Sciencal, Carlsbad, CA) were cultured in DMEM containing 1% penicillin/streptomycin, L-glutamine and 10% fetal bovine serum (Atlanta biologics, Flowery Branch, GA). NHLFs derived from 3 separate donors were utilized for experiments. Cells were passaged upon achieving 90% confluency at a 1:4 ratio and used for studies until passage 7. For all hydrogel studies, cells were trypsinized, counted and either encapsulated into or seeded onto 25 µl hydrogels at a density of 1,000,000 cells ml^-1^ of hydrogel, and subsequently cultured at 37°C and 5% CO2 in serum containing medium. For studies comparing 3D hydrogels to tissue culture plastic, the number of cells seeded into 2D conditions was analogous to the total cell number within hydrogel matrices. Media was refreshed the day after encapsulation and every 2 days after. In selected experiments, recombinant human TGF-β1 (5 ng/ml, Peprotech, Rocky Hill, NJ) was supplemented into the media at 5 ng ml^-1^. For pharmacological studies, Nintedanib (50 nM, Fisher Scientific, Hampton, NH), Pirfenidone (100 µM, Fisher Scientific, Hampton, NH), Marimastat (1.0 µM), DMF (100 nM), were supplemented in serum containing media and refreshed every 2 days.

### Fluorescent staining, microscopy, and analysis

Cultures were fixed with 4% paraformaldehyde for 30 min at room temperature. To stain the actin cytoskeleton and nuclei, samples were permeabilized in PBS solution containing Triton X-100 (5% v/v), sucrose (10% w/v), and magnesium chloride (0.6% w/v), blocked in 1% BSA, and stained simultaneously with phalloidin and DAPI. For immunostaining, samples were permeabilized, blocked for 8 h in 1% w/v bovine serum albumin, and incubated with mouse monoclonal anti-YAP antibody (1:1000, Santa Cruz SC-101199), mouse monoclonal anti-fibronectin antibody (FN, 1:2000, Sigma #F6140), rabbit monoclonal anti-Ki67 (1:500, Sigma #PIMA514520) or mouse monoclonal anti-α-SMA (1:2000, Sigma #A2547) followed by secondary antibody for 6 h each at room temperature with 3x PBS washes in between. High-resolution images of YAP, FN and actin morphology were acquired with a 40x objective. Unless otherwise specified, images are presented as maximum intensity projections of 100µm Z-stacks. Hydrogel samples were imaged on a Zeiss LSM 800 laser scanning confocal microscope. SHG imaging of lung tissue was imaged was conducted on a Leica SPX8 laser scanning confocal microscope with an excitation wavelength of 820 nm and a collection window of 400-440 nm. Single cell morphometric analyses (cell spread area) were performed using custom Matlab scripts with sample sizes > 50 cells/group, while YAP, α-SMA, Ki67, FN immunostains were quantified on an image basis with a total of 10 frames of view. For cell density (# of nuclei) calculations, DAPI-stained cell nuclei were thresholded and counted in six separate 600 x 600 x 200 µm image volumes, allowing us to calculate a total number of cells per mm ^3^ of gel. Fiber recruitment analysis was conducted via a custom Matlab script; briefly, cell outlines were created via actin masking and total fiber fluorescence was quantified under each actin mask on a per cell basis.

### RT-qPCR

For all experiments, additional hydrogel replicates were finely minced and degraded in dextranase solution (4 IU/ml, Sigma) for 20 minutes, homogenized in buffer RLT (Qiagen, Venlo, Netherlands), and RNA was isolated according to manufacturer protocols. cDNA was generated from DNAse-free RNA, amplified, and gene expression was normalized to the house-keeping gene, glyceraldehyde 3-phosphate dehydrogenase (GAPDH). Experiments were run with technical triplicates across three individual biological experiments. For a complete list of primers, see Supplementary Table S2.

### Mechanical testing

To determine the elastic modulus of lung tissue and DexVS hydrogels, indentation tests were employed using a Nanosurf FlexBio atomic force microscope (AFM; Nanosurf, Liestal, Switzerland). Samples were indented upon via a 5 µm bead at a depth of 10 µm and an indentation rate of 0.333 µm/s. Resulting force-displacement curves were fit to a spherical Hertz model utilizing AtomicJ. A Poisson’s ratio of 0.5 and 0.4 were utilized for hydrogels and lung tissue, respectively.

### Animal studies

All animal studies were approved by the Animal Care and Use Committee at the University of Michigan. Bleomycin (0.025 U; Sigma) was instilled intratracheally in C57BL6 mice (8 wk of age; The Jackson Laboratory, Bar Harbor, ME, USA) on day 0. Briefly, mice were anesthetized with sodium pentobarbital, the trachea was exposed and entered with a 30-gauge needle under direct visualization, and a single 30-μl injection containing 0.025 U bleomycin (Sigma) diluted in normal saline was injected. Lungs were collected on day 14 for mechanical and histological analysis. For histology samples, lungs were perfused with saline and inflated with 4% paraformaldehyde, sectioned, and stained with Picosirius Red. For mechanical characterization via AFM, lungs were perfused with saline, infused with OCT compound (Fisher), and flash frozen in a slurry of dry ice and ethanol. Sections were mechanically tested via AFM nanoindentation immediately upon thawing.

### Cytokine and collagen measurements

To characterize the inflammatory secretome associated with various DexVS-VPMS environments, media was collected from NHLF cultures 3, 5, 7, and 9 days post encapsulation. A Luminex FlexMAP 3D (Luminex Corporation, Austin, TX) systems technology was used to measure 41 cytokines/ chemokines (HCTYMAG-60K-PX41, Milliplex, EMD Millipore Corporation) in the media samples according to manufacturer instructions. Total secretion was reported as the sum of all 41 analytes measured for each respective condition. Cell secreted collagen was measured using the established colorimetric Sircol assay in hydrogels cultured with serum free media in the presence of 25 µg ml^-1^ ascorbic acid.

### Bioinformatics

The NCBI GEO database was consulted (dataset GSE47460 (GP14550)) to fetch gene expression data from 91 healthy patients and 140 patients with IPF; patients with COPD and non-idiopathic fibrotic lung diseases were excluded from the analysis (*44*). GEO2R software was utilized for GO term enrichment, with keywords extracellular matrix, MMP, integrin, cytoskeleton, cytokine, chemokine, and MAPK used as search terms for dataset curation. Noncurated datasets were composed of the top 1000 differentially expressed genes between healthy and ILD conditions. For pathway and protein-protein enrichment analyses, a curated pathway database (Reactome (*45*)) and Search Tool for Retrieval of Interacting Genes/Proteins (STRING (*46*)) methodology were consulted, respectively. For STRING analyses, upregulated genes within the druggable genome were focused upon. A tabulated list of top genes, pathways, and nodes can be seen in Supplementary Data File S1.

### Statistical Analysis

Statistical significance was determined by one-way analysis of variance (ANOVA) or Student’s t-test where appropriate, with significance indicated by p<0.05. All data are presented as mean ± standard deviation.

## General

We thank Dr. Eric S. White (University of Michigan) for providing patient-derived lung fibroblasts employed in these studies.

## Funding

This work was supported in part by the National Institutes of Health (HL124322). DLM acknowledges financial support from the University of Michigan Rackham Merit Fellowship and the National Science Foundation Graduate Research Fellowship Program (DGE1256260).

## Author Contributions

D.L.M and B.M.B conceived and supervised the project. D.L.M. designed and performed the experiments. K.M.D and K.B.A performed and aided in analysis of the Luminex experiments. M.R.S and C.D.D. helped with data analysis. R.P. and M.S. aided in polymer syntheses and microfiber fabrication. I.M.L provided equipment for and assisted in PCR experiments. C.A.W and B.B.M helped perform the animal experiments for the Bleomycin-induced lung fibrosis model. All authors edited and approved the manuscript.

## Competing interest

Authors disclose no competing or financial conflicts of interest.

## Data and materials availability

All additional data needed to evaluate the conclusions in this paper are present in the paper and/or the Supplementary Materials. Additional data related to this paper may be requested from the D.L.M and B.M.B.

## Supplementary Information

**Figure S1:**
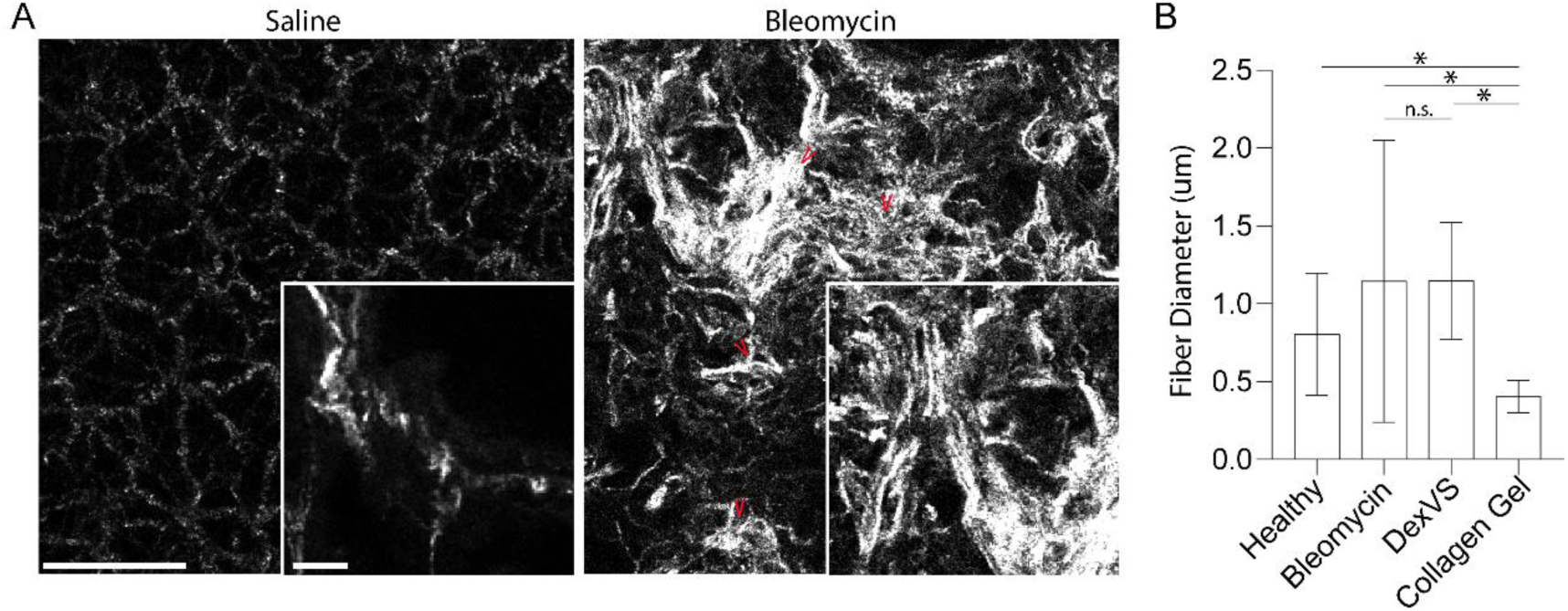
Fiber structure within saline and bleomycin treated lungs. A) Second harmonic generation microscopy images of collagen fibers within saline and bleomycin treated mouse lung tissue (scale bar: 75 µm); insets: higher resolution images depicting regions containing collagen fiber bundles in bleomycin treated lung tissue (scale bar: 5 µm). B) Quantification of fiber diameter (n>20 fibers) within mouse lung compared to synthetic DexVS fibers and collagen fibers within 3mg/ml reconstituted type I collagen matrices.

**Figure S2:**
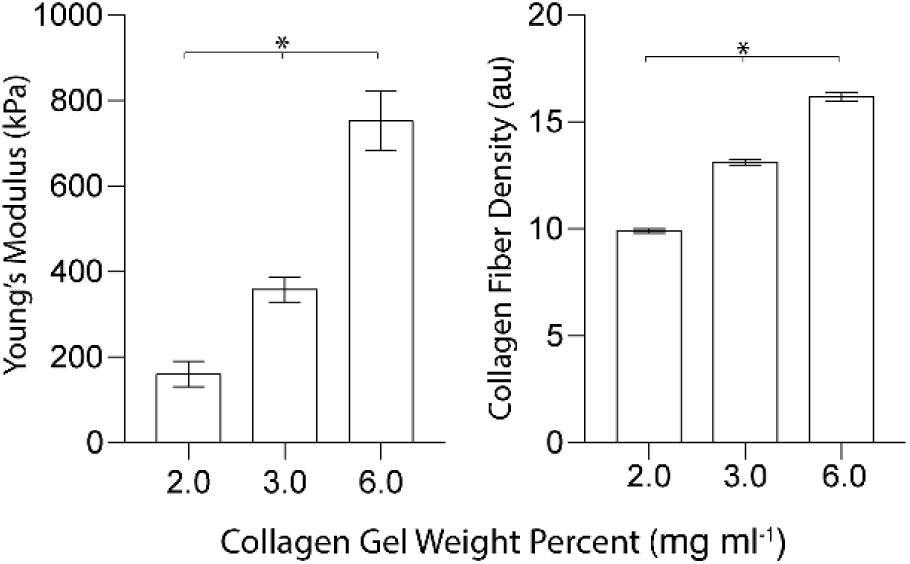
Relationship between collagen gel mechanical and structural properties. Quantification of type I collagen gel Young’s modulus and fiber density as a function of gel weight percentage (n=10 measurements/group).

**Figure S3:**
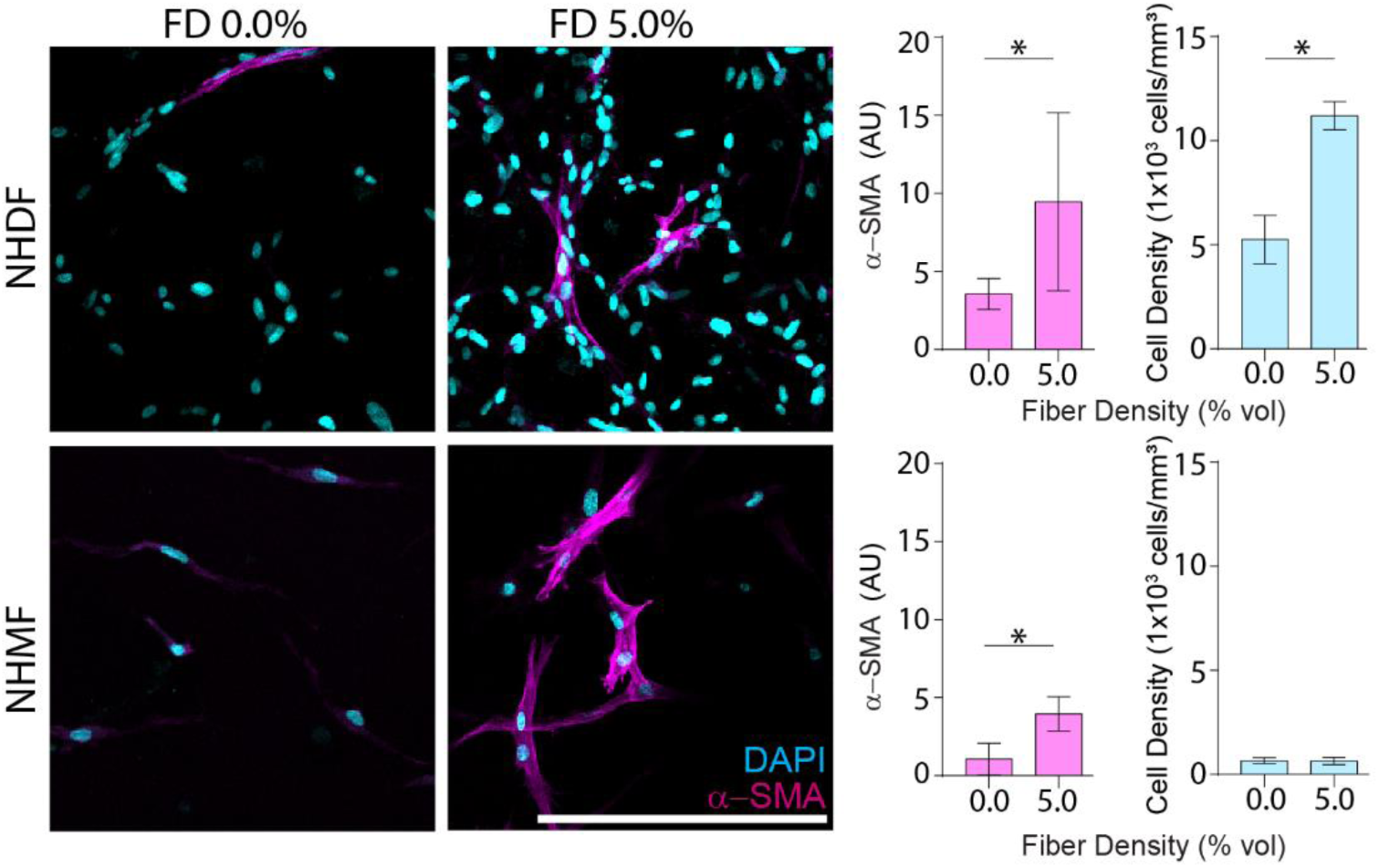
Fiber density effects on dermal and mammary fibroblasts. Representative images of NHDF and NHMF within nonfibrous control and fibrous hydrogels after 9 days of culture (scale bar: 200 µm), with quantification of α-SMA and cell density (n=4 samples/group with n=10 frames of view quantified and n>25 cells/frame of view).

**Figure S4:**
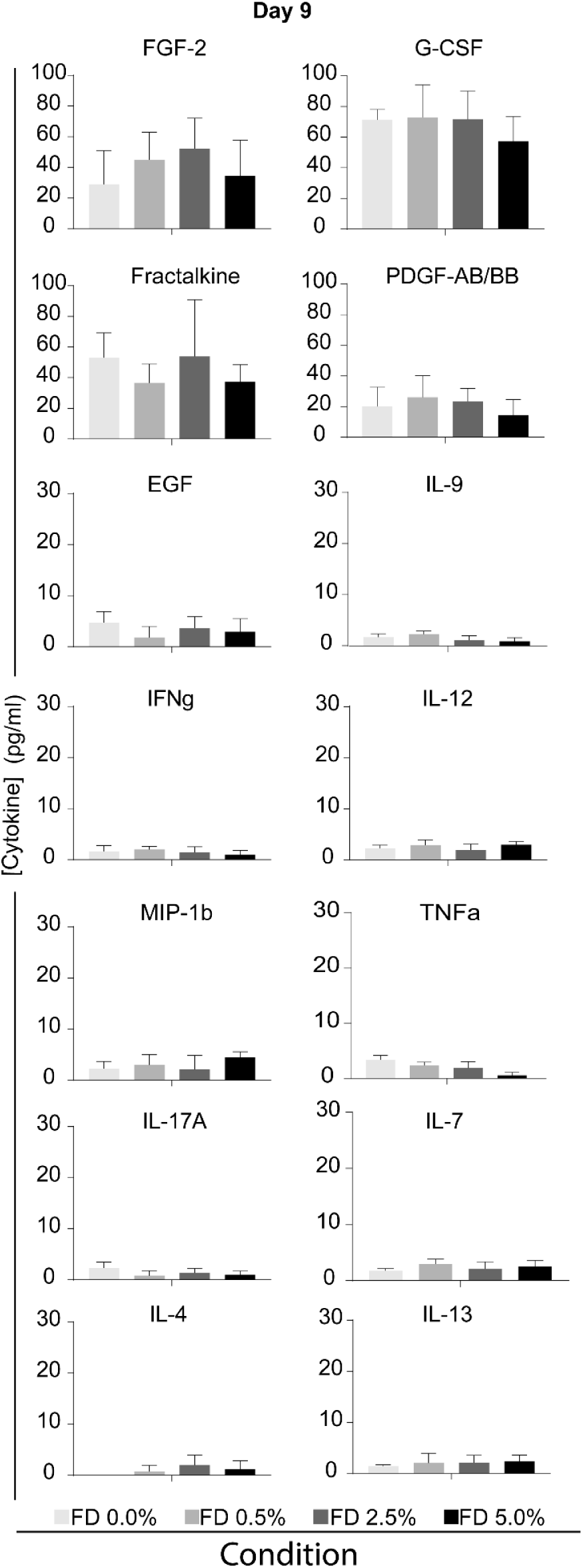
Measurable cytokines in 3D fibroblast supernatant. Luminex quantification of detectable cytokines and growth factors in day 9 NHLF culture media supernatant over a range of fiber density (n=4 samples/group). Factors which significantly changed as a function of fiber density can be seen in Figure 4.

**Table S1:** Example matrix remodeling genes which are associated with IPF.

| **Example Matrix**<br>**Remodeling Genes** | **Differential**<br>**Expression**<br>**(IPF - Healthy)** | **adj.P.Val** |
| --- | --- | --- |
| MMP1 | 4.623214176 | 5E-31 |
| MMP7 | 2.975102692 | 2E-30 |
| MMP9 | 0.420781319 | 6E-02 |
| MMP10 | 2.664035989 | 1E-15 |
| MMP11 | 2.224064505 | 7E-28 |
| MMP3 | 1.682079176 | 3E-16 |
| MMP12 | 1.122249176 | 2E-04 |
| COL1A2 | 0.950009835 | 1E-23 |
| LOX | 0.429231923 | 1.9E-09 |
| ACAN | 1.365209286 | 9.61E-14 |
| DCN | 0.171328901 | 0.006698 |
| HS6ST3 | 1.949643791 | 1.22E-20 |
| HS6ST2 | 2.817251484 | 9.46E-43 |

**Table S2:**
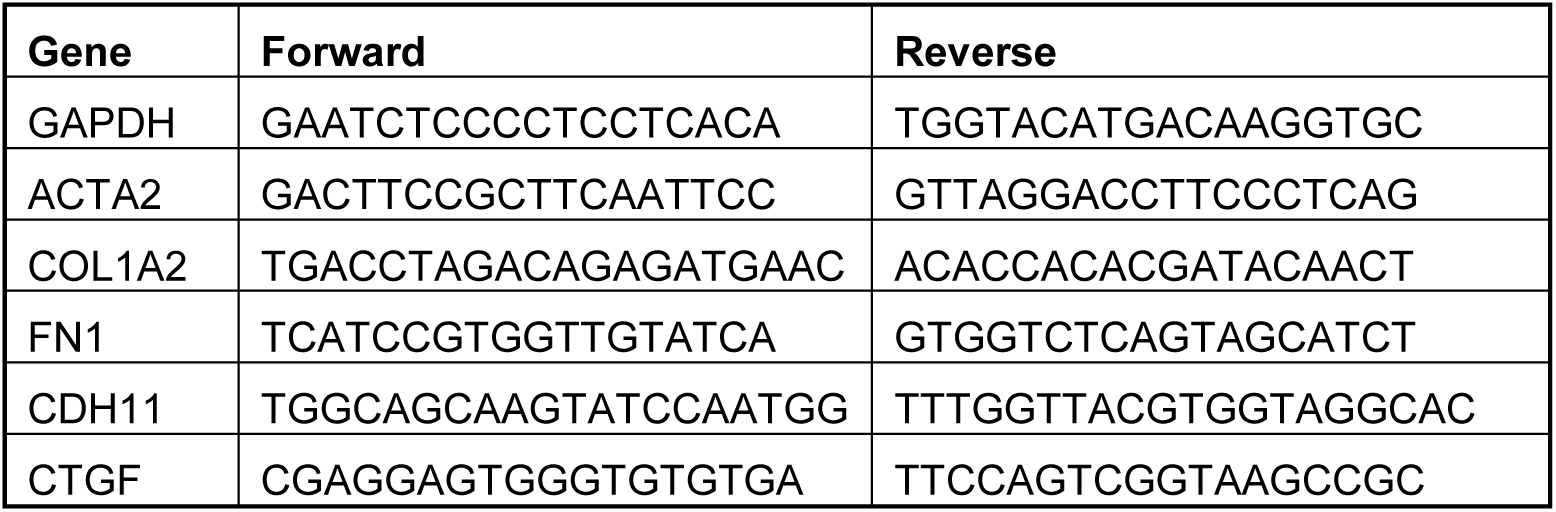
Primer list for genes associated with MF differentiation (Figure 3).

## Notes

### Competing Interest Statement

The authors have declared no competing interest.

## References

1. T. A. Wynn, Cellular and molecular mechanisms of fibrosis. J. Pathol. 214, 199–210 (2008).

2. G. Bagnato, S. Harari, Cellular interactions in the pathogenesis of interstitial lung diseases. Eur. Respir. Rev. 24, 102–114 (2015).

3. V. F. Fiore, S. Wong, C. Tran, C. Tan, W. Xu, E. S. White, J. S. Hagood, T. H. Barker, Integrin a v b 3 drives fibroblast contraction and strain stiffening of soft provisional extracellular matrix during progressive fibrosis. 3, 1–35 (2018).

4. A. Sundarakrishnan, Y. Chen, L. D. Black, B. B. Aldridge, D. L. Kaplan, Engineered cell and tissue models of pulmonary fibrosis. Adv. Drug Deliv. Rev. 129, 78–94 (2018).

5. A. Pathak, S. Kumar, Biophysical regulation of tumor cell invasion: moving beyond matrix stiffness. Integr Biol. 3, 267–278 (2011).

6. P. D. Arora, N. Narani, C. A. G. McCulloch, The Compliance of Collagen Gels Regulates Transforming Growth Factor-β Induction of α-Smooth Muscle Actin in Fibroblasts. Am. J. Pathol. 154, 871–882 (1999).

7. S. Nakagawa, P. Pawelek, F. Grinnell, Long-Term Culture of Fibroblasts in Contracted Collagen Gels: Effects on Cell Growth and Biosynthetic Activity. J. Invest. Dermatol. 93, 792–798 (1989).

8. M. R. Ebrahimkhani, C. L. Young, D. A. Lauffenburger, L. G. Griffith, J. T. Borenstein, Approaches to in vitro tissue regeneration with application for human disease modeling and drug development. Drug Discov. Today. 19, 754–762 (2014).

9. L. F. Castella, L. Buscemi, C. Godbout, J. J. Meister, B. Hinz, A new lock-step mechanism of matrix remodelling based on subcellular contractile events. J. Cell Sci. 123, 1751–1760 (2010).

10. J. L. Balestrini, S. Chaudhry, V. Sarrazy, A. Koehler, B. Hinz, The mechanical memory of lung myofibroblasts. Integr. Biol. 4, 410–421 (2012).

11. M.-L. Bochaton Piallat, G. Gabbiani, B. Hinz, The Myofibroblast in wound healing and fibrosis: Answered and unanswered questions. F1000 Res. 5 (2016), doi: 10.12688/f1000research.8190.1.

12. A. Giménez, P. Duch, M. Puig, M. Gabasa, A. Xaubet, J. Alcaraz, Dysregulated collagen homeostasis by matrix stiffening and TGF-β1 in fibroblasts from idiopathic pulmonary fibrosis patients: Role of FAK/Akt. Int. J. Mol. Sci. 18 (2017), doi: 10.3390/ijms18112431.

13. F. Liu, D. J. Tschumperlin, Micro-mechanical characterization of lung tissue using atomic force microscopy. J. Vis. Exp., 1–7 (2011).

14. P. Zigrino, J. Brinckmann, A. Niehoff, Y. Lu, N. Giebeler, B. Eckes, K. E. Kadler, C. Mauch, Fibroblast-Derived MMP-14 Regulates Collagen Homeostasis in Adult Skin. J. Invest. Dermatol. 136, 1575–1583 (2016).

15. M. Corbel, J. Lanchou, N. Germain, Y. Malledant, E. Boichot, V. Lagente, Modulation of airway remodeling-associated mediators by the antifibrotic compound, pirfenidone, and the matrix metalloproteinase inhibitor, batimastat, during acute lung injury in mice. Eur. J. Pharmacol. 426, 113–121 (2001).

16. S. R. Caliari, S. L. Vega, M. Kwon, E. M. Soulas, J. A. Burdick, Dimensionality and spreading influence MSC YAP/TAZ signaling in hydrogel environments. Biomaterials. 103, 314–323 (2016).

17. L. Follonier, S. Schaub, J.-J. Meister, B. Hinz, Myofibroblast communication is controlled by intercellular mechanical coupling. J. Cell Sci. 121, 3305–3316 (2008).

18. R. M. Kottmann, J. Sharp, K. Owens, P. Salzman, G. Q. Xiao, R. P. Phipps, P. J. Sime, E. B. Brown, S. W. Perry, Second harmonic generation microscopy reveals altered collagen microstructure in usual interstitial pneumonia versus healthy lung. Respir. Res. 16, 1–13 (2015).

19. D. L. Matera, W. Y. Wang, M. R. Smith, A. Shikanov, B. M. Baker, Fiber Density Modulates Cell Spreading in 3D Interstitial Matrix Mimetics (2019), doi: 10.1021/acsbiomaterials.9b00141.

20. F. Liu, D. Lagares, K. M. Choi, L. Stopfer, A. Marinković, V. Vrbanac, C. K. Probst, S. E. Hiemer, T. H. Sisson, J. C. Horowitz, I. O. Rosas, L. E. Fredenburgh, C. Feghali-Bostwick, X. Varelas, A. M. Tager, D. J. Tschumperlin, Mechanosignaling through YAP and TAZ drives fibroblast activation and fibrosis. Am. J. Physiol. - Lung Cell. Mol. Physiol. 308 (2015).

21. I. G. Luzina, N. W. Todd, S. Sundararajan, S. P. Atamas, The cytokines of pulmonary fibrosis: Much learned, much more to learn. Cytokine. 74, 88–100 (2015).

22. M. Gharaee-kermani, R. E. Mccullumsmith, I. F. Charo, S. L. Kunkel, S. H. Phan, CC-chemokine receptor 2 required for bleomycin-induced pulmonary fibrosis. 24, 266–276 (2003).

23. H. Sahin, H. E. Wasmuth, Chemokines in tissue fibrosis. Biochim. Biophys. Acta - Mol. Basis Dis. 1832, 1041–1048 (2013).

24. B. Hinz, Formation and Function of the Myofibroblast during Tissue Repair. J. Invest. Dermatol. 127, 526–537 (2007).

25. M. Carone, F. G. Salerno, A. M. Esquinas, Obstructive lung function decline and IPF. Chron. Respir. Dis. 13, 204–205 (2016).

26. T. Ogura, H. Taniguchi, A. Azuma, Y. Inoue, Y. Kondoh, Y. Hasegawa, M. Bando, S. Abe, Y. Mochizuki, K. Chida, M. Klüglich, T. Fujimoto, Safety and pharmacokinetics of nintedanib and pirfenidone in idiopathic pulmonary fibrosis, 1382–1392.

27. A. P. Grzegorzewska, F. Seta, R. Han, C. A. Czajka, K. Makino, L. Stawski, J. S. Isenberg, J. L. Browning, M. Trojanowska, Dimethyl Fumarate ameliorates pulmonary arterial hypertension and lung fibrosis by targeting multiple pathways. Sci. Rep. 7, 1–14 (2017).

28. V. E. de Meijer, D. Y. Sverdlov, Y. Popov, H. D. Le, J. A. Meisel, V. Nosé, D. Schuppan, M. Puder, Broad-spectrum matrix metalloproteinase inhibition curbs inflammation and liver injury but aggravates experimental liver fibrosis in mice. PLoS One. 5 (2010), doi: 10.1371/journal.pone.0011256.

29. C. M. Madl, B. L. Lesavage, R. E. Dewi, C. B. Dinh, R. S. Stowers, M. Khariton, K. J. Lampe, D. Nguyen, O. Chaudhuri, A. Enejder, S. C. Heilshorn, Maintenance of neural progenitor cell stemness in 3D hydrogels requires matrix remodelling. Nat. Mater. 16, 1233–1242 (2017).

30. E. W. Howard, B. J. Crider, D. L. Updike, E. C. Bullen, E. E. Parks, C. J. Haaksma, D. M. Sherry, J. J. Tomasek, MMP-2 expression by fibroblasts is suppressed by the myofibroblast phenotype. Exp. Cell Res. 318, 1542–1553 (2012).

31. M. Ohshima, Y. Yamaguchi, K. Ambe, M. Horie, A. Saito, T. Nagase, K. Nakashima, H. Ohki, T. Kawai, Y. Abiko, P. Micke, K. Kappert, Fibroblast VEGF-receptor 1 expression as molecular target in periodontitis. J. Clin. Periodontol. 43, 128–137 (2016).

32. K. P. Sundararaj, D. J. Samuvel, Y. Li, J. J. Sanders, M. F. Lopes-Virella, Y. Huang, Interleukin-6 released from fibroblasts is essential for up-regulation of matrix metalloproteinase-1 expression by U937 macrophages in coculture: Cross-talking between fibroblasts and U937 macrophages exposed to high glucose. J. Biol. Chem. 284, 13714–13724 (2009).

33. M. M. Mia, M. Boersema, R. A. Bank, Interleukin-1β attenuates myofibroblast formation and extracellular matrix production in dermal and lung fibroblasts exposed to transforming growth factor-β1. PLoS One. 9 (2014), doi: 10.1371/journal.pone.0091559.

34. M. R. M. H. Bart van der Worp, D. W. Howells, E.S. Sena, M.J. Porritt, S. Rewell, V. O’Collins, Can Animal Models of Disease Reliably Inform Human Studies? PLoS One. 11 (2016), doi: 10.1371/journal.

35. T. R. Cox, J. T. Erler, Remodeling and homeostasis of the extracellular matrix: implications for fibrotic diseases and cancer. Dis. Model. Mech. 4, 165–178 (2011).

36. A. Santos, D. Lagares, Matrix Stiffness : the Conductor of Organ Fibrosis (2018).

37. G. Raghu, K. K. Brown, H. R. Collard, V. Cottin, K. F. Gibson, R. J. Kaner, D. J. Lederer, F. J. Martinez, P. W. Noble, J. W. Song, A. U. Wells, T. P. M. Whelan, W. Wuyts, E. Moreau, S. D. Patterson, V. Smith, S. Bayly, J. W. Chien, Q. Gong, J. J. Zhang, T. G. O’Riordan, Efficacy of simtuzumab versus placebo in patients with idiopathic pulmonary fibrosis: a randomised, double-blind, controlled, phase 2 trial. Lancet Respir. Med. 5, 22–32 (2017).

38. S. Piera-Velazquez, Z. Li, S. A. Jimenez, Role of Endothelial-Mesenchymal Transition (EndoMT) in the Pathogenesis of Fibrotic Disorders. Am. J. Pathol. 179, 1074–1080 (2011).

39. J. Ehling, M. Bartneck, X. Wei, F. Gremse, V. Fech, D. Möckel, C. Baeck, K. Hittatiya, D. Eulberg, T. Luedde, F. Kiessling, C. Trautwein, T. Lammers, F. Tacke, CCL2-dependent infiltrating macrophages promote angiogenesis in progressive liver fibrosis. Gut. 63, 1960–1971 (2014).

40. M. Asmani, S. Velumani, Y. Li, N. Wawrzyniak, I. Hsia, Z. Chen, B. Hinz, R. Zhao, Fibrotic microtissue array to predict anti-fibrosis drug efficacy. Nat. Commun. 9, 1–12 (2018).

41. Y. Kurita, J. Araya, S. Minagawa, H. Hara, A. Ichikawa, N. Saito, T. Kadota, K. Tsubouchi, N. Sato, M. Yoshida, K. Kobayashi, S. Ito, Y. Fujita, H. Utsumi, H. Yanagisawa, M. Hashimoto, H. Wakui, Y. Yoshii, T. Ishikawa, T. Numata, Y. Kaneko, H. Asano, M. Yamashita, M. Odaka, T. Morikawa, Pirfenidone inhibits myofibroblast differentiation and lung fibrosis development during insufficient mitophagy, 1–14 (2017).

42. J. Schuett, A. Ostermann, L. Wollin, The Effect Of Nintedanib Compared To Pirfenidone On Serum-Stimulated Proliferation Of Human Primary Lung Fibroblasts At Clinically Relevant Concentrations. Am J Respir rit Care Med. 191, A4940 (2015).

43. B. M. Baker, B. Trappmann, W. Y. Wang, M. S. Sakar, I. L. Kim, V. B. Shenoy, J. A. Burdick, C. S. Chen, Cell-mediated fibre recruitment drives extracellular matrix mechanosensing in engineered fibrillar microenvironments. Nat. Mater. 14, 1262–1268 (2015).

44. T. Barrett, S. E. Wilhite, P. Ledoux, C. Evangelista, I. F. Kim, M. Tomashevsky, K. A. Marshall, K. H. Phillippy, P. M. Sherman, M. Holko, A. Yefanov, H. Lee, N. Zhang, C. L. Robertson, N. Serova, S. Davis, A. Soboleva, NCBI GEO: Archive for functional genomics data sets - Update. Nucleic Acids Res. 41, 991–995 (2013).

45. B. Jassal, L. Matthews, G. Viteri, C. Gong, P. Lorente, A. Fabregat, K. Sidiropoulos, J. Cook, M. Gillespie, R. Haw, F. Loney, B. May, M. Milacic, K. Rothfels, C. Sevilla, V. Shamovsky, S. Shorser, T. Varusai, J. Weiser, G. Wu, L. Stein, H. Hermjakob, P. D’Eustachio, The reactome pathway knowledgebase. Nucleic Acids Res., 2–7 (2019).

46. D. Szklarczyk, A. L. Gable, D. Lyon, A. Junge, S. Wyder, J. Huerta-Cepas, M. Simonovic, N. T. Doncheva, J. H. Morris, P. Bork, L. J. Jensen, C. Von Mering, STRING v11: Protein-protein association networks with increased coverage, supporting functional discovery in genome-wide experimental datasets. Nucleic Acids Res. 47, D607–D613 (2019).

